# Color appearance of iridescent objects

**DOI:** 10.1101/2023.04.12.535824

**Authors:** Katja Doerschner, Robert Ennis, Philipp Börner, Frank J. Maile, Karl R. Gegenfurtner

## Abstract

Iridescent objects and animals are quite mesmerizing to look at, since they feature multiple intense colors whose distribution can vary quite dramatically as a function of viewing angle. These properties make them a particularly interesting and unique stimulus to experimentally investigate the factors that contribute to single color impressions of multi-colored objects. Our stimuli were 3D printed shapes of varying complexity that were covered with three different types of iridescent coatings. For each shape-color combination, participants performed single- and multi-color matches for different views of the stationary object, as well as single color matches for a corresponding rotating stimulus. In the multi-color matching task, participants subsequently rated the size of the surface area on the object that was covered by the match-identified color. Results show that single-color appearance of iridescent objects varied with shape complexity, view, and object motion. Moreover, hue similarity of color settings in the multi color match task best predicted single color appearance, however this predictor was weaker for predicting single color matches in the motion condition. Taken together our findings suggest that the single color appearance of iridescent objects may be modulated by chromatic factors, spatial-relations and the characteristic dynamics of color changes that are typical for this type of material.

## 1 Introduction

Iridescence refers to the phenomenon that the color of an object changes with viewing angle. It can occur for example, on bird feathers, fish scales or beetles and is believed to have some important biological functions [1][2][3]. Also many non-living things like soap bubbles, oily films on water, or mother of pearl have iridescent qualities. Iridescence is an appealing optical phenomenon that can have multiple physical causes and to understand, model and produce iridescent materials is an active area of research [4][5][6][7][8].

Describing the color appearance of an iridescent object is challenging with a single color term, for the above reason (also see a nice early treatment of this in [9]). Consider, for example, the iridescent objects in Fig.1A. Each column shows one type of iridescent paint that each features about 2-3 prominent colors. How would one go about deciding the dominant color for each one of these? Is the object in Fig1. A.1 purple with some green or vice versa? How about the object in Fig.1 A.4? What does this decision depend on? The saturation or salience of a color? The hue similarity of colors? Or more geometrical aspects e.g. how much area on the object is taken up by a color, or whether a color is located towards the object boundary or the center? A decision based on geometrical factors is likely to get complicated, because as the shape of the objects gets more undulated (rows), the spatial distribution of colors on the object becomes more complex, since the ‘occurrence’ of a given shade of color is dependent on the angle between surface normal and line of sight. While our little thought experiment highlights the challenges that one might face when color categorizing iridescent objects, it is precisely these properties that make them a particularly interesting class of stimuli to study color appearance and categorization of multicolored objects per se, because they allow us to test several specific geometric and colorimetric hypotheses in single object color perception.

Even for non-iridescent stimuli, forming a single color impression of multicolored objects is a potentially complicated process, as the color impression not only requires invariant estimates of hue, saturation, and brightness of the object’s color, as would be the case for uniformly colored objects (e.g. as in [10]), but also requires some type of additional process that integrates multiple colors into a single color impression.

One such process could be photometric averaging. Using random dot stimuli consisting of two shades of color with the same hue and brightness, but different saturation, [11] tested this idea and found that the visual system does not rely on a photometric average color to form a single color impression of a multi-coloured stimulus. Rather, it integrates the color appearances (perceived hue, saturation, brightness) of individual colors to form a single color impression. Moreover, the single color impression was shifted to the more saturated color. In a similar experiment, [12] found that the greater the range of colors in a mosaic pattern the more participants’ color matches deviated from the photometric average of a given stimulus, showing that the element with the highest-saturation strongly influenced the single color impression and that photometric averaging is less likely to occur for dissimilar hues. These studies provide insights into when photometric averaging might come into play when forming single color estimates of multicolored stimuli. However, it is not clear how well these results that were obtained with simple random dot and checker patterns will generalize to more natural objects, that often exhibit larger patches of varying color or that have smooth transitions between different color patches.

[13] used such natural stimuli, asking participants to classify fall leaves as red or green. The leaves were shown as photographs and exhibited red and green patches to various amounts. The authors found that participants’ responses could be well predicted by the image-based average hue angle of a given leaf (regardless of saturation and brightness), with the individual participant’s unique yellow hue angle serving as a boundary between red and green, and they concluded that participants based their decision on an extraction of an average hue. In a second experiment, the authors “flipped” the leaves’ color distribution, by mirroring each individual pixel’s hue angle about the average hue angle of the whole leaf, while keeping the saturation and brightness constant. The average hue angle was unchanged through this process. If participants relied on an average hue angle to determine the single color impression of a given leaf, then one would assume that their categorizations should remain the same. For most stimuli, this was the case, but for some leaves, there were changes in categorization consistent across participants. While the authors hold on to their general interpretation, namely, that participants extract an average hue angle and compare it to their internal decision boundary of unique yellow to form their decision, they also acknowledge that for those cases where the classification changed for flipped stimuli, there must be additional factors, that could have an effect on the decision in addition to the average hue. Potential explanations for the change in categorization for stimuli near the unique yellow decision boundary might have been subtle changes in apparent saturation, or changes in the spatial distribution of red and green colors brought on by the pixel value flip. In fact, inspecting their Fig. 12 [13] it seems like the color that the leaf was more often categorized as, was also the color that occupied more of the leaf’s surface.

Taken together, results from the reviewed literature suggest that single color estimation of multicolored objects is a complex process that is modulated by several factors, including chromatic properties, e.g. saturation or hue similarity of colors, and geometric aspects, e.g. the amount and location of a color patch on a surface. However, much research still needs to be done to fully map out and understand the complex interplay of these factors.

In the experiments described below, we use the well established color matching paradigm by [10] together with a novel type of stimulus, namely manufactured, iridescent objects of varying shape complexity (Fig. 1A) to advance our knowledge on the formation of single color impressions of multicolored objects. The iridescent paints feature color combinations with similar (Fig. 1A, column 2) and dissimilar hues (Fig. 1A, column 1), as well as combinations with more and less saturated hues (Fig. 1A, column 3, for a less saturated example), which allows us to test to what extent hue similarity and saturation predict the overall color appearance of these objects. Moreover, the complex relationship between the occurrence of a color, object geometry and viewing angle not only lets us investigate whether the total area, or coverage, of a color on the object predicts overall color appearance but also more specific questions, i.e. to what extent the location of a color patch modulates appearance, e.g. whether a color patch occupies central or near-contour regions on the object. Finally, color changes on iridescent objects are especially dramatic as the object moves, or as we move with respect to the object. This feature opens up unprecedented possibilities to investigate how object motion and associated changes in appearance over time modulate single color impressions. Details of how we test each of these questions are described next.

**Figure 1.**
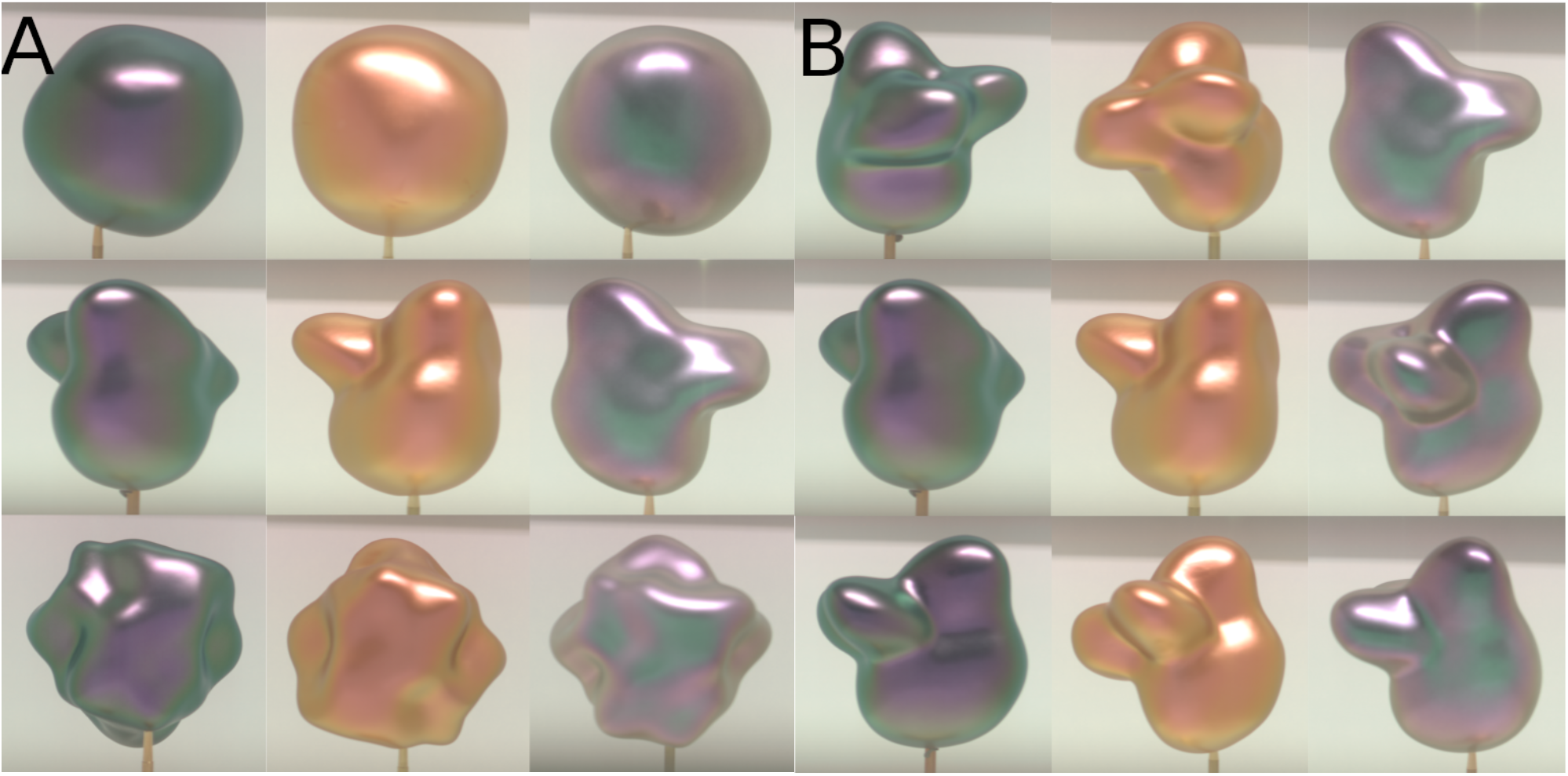
Stimuli. Hyperspectral photos (VNIR HS-CL-30-VSE-OEM mirror camera from Specim (Specim, Spectral Imaging, Ltd.; Oulu, Finnland)) of the three shapes (rows) coated each with three types of goniochromatic paints (provided by Schlenk Metallic Pigments GmbH; Roth, Germany). Shapes were approximately 14 cm in diameter which corresponds to about 16.5 deg visual angle. Viewing angles were chosen to maximize appearance variation. Panel A shows stimuli organized according to shape complexity (rows) from lowest (shape 1) to highest (shape 3). Panel B shows shape 2 in each paint, at three possible views, as an example to illustrate the marked changes in appearance that can occur across viewing positions. These changes are most pronounced at intermediate levels of shape complexity. Note that the views chosen differed slightly between the differently painted objects. This was due to the fact that the insertion point for the metal stand was not identical between same-shaped objects. This variation is not relevant to any of our results or the overall conclusion.

## 2 Methods

### Participants, Stimuli & Apparatus

Seven voluntary participants with normal or corrected-to-normal vision participated in the experiments, and were reimbursed with 8€/h for their efforts. All experiments were approved by the local ethics review board and conducted in accordance with the Declaration of Helsinki.

Three different shapes of three levels of complexity each [14][15] were 3D-printed. To generate the goniochromatic color stimuli, the objects were modified with different coatings containing silica flake-based effect pigments (see [6] for details, and Fig. 1A). These shape-color combinations created a unique color appearance for each stimulus that could vary profoundly as a function of viewing angle (Fig. 1B). Objects were illuminated by standard office neon tube lighting and were mounted on a metal stand that was 16 cm in height and placed inside a large white box (78 × 59 × 53 cm), with an opening at the top, a viewing slit (15 × 3.5 cm) for the participant at one side and a door for the experimenter (for easy exchange of the stimuli) at the opposite side. Stimuli were either placed on a pedestal or a rotating disc (counterclockwise rotation at 2rpm), both 6 cm in height and approximately 48 cm away from the viewing slit. Participants performed appearance tasks on a calibrated computer monitor (ColorEdge CG223W; EIZO Corporation; Hakusan, Japan) that was positioned next to the white box, so that participants could easily change their gaze between stimulus and monitor. The computer running the experiment was a Dell Precision 390 Desktop (Dell, Inc.; Round Rock, Texas, USA) using Windows 7 Professional (64-bit) as operating system with Service Pack 1 (Microsoft Corporation; Redmond, Washington, USA) with a NVidia Quadro 2000 graphics card (Nvidia Corporation; Santa Clara, California, USA). Experimental programs were written in MATLAB (R2007b) using Psychtoolbox (SVN Revision 4881, cloned from Github) [16][17][18].

### Procedure

The experiment was divided into a static and a motion block. Participants completed three tasks in the first and one task in the second block. In the static block, on every trial, the experimenter placed a randomly chosen (via computer program) stimulus on the pedestal in the presentation box. Note, that objects of the same paint were never shown on consecutive trials. Participants were instructed to not look through the viewing slit during the placement to prevent them from seeing the object in motion and at different viewing angles. The first task on a given trial was to choose a single color to describe the object, by adjusting the hue, saturation and lightness of a circular test patch presented on the monitor (single match task). To do this, the participant used a computer mouse to navigate through the DKL color space. Participants could lock the color by clicking space and they pressed “F” and “Enter”on a keyboard to confirm their answer. For the second task on a given trial, participants could color-adjust two or three circular patches to indicate multiple colors that they perceived to be most prominent on the object (multi-match task). Participants indicated, for example, ‘2’, if they perceived only two prominent colors. In that case they would only be presented with two disks for the color adjustment task. Following this, in the third task, participants saw the results of color adjustments of task 2 and, by moving sliders, indicated the relative area on the object that was covered by each of the colors. The sliders were not yoked so it was technically possible to give answers that do not add up to 100%. During a trial, participants were allowed to look at the object as often and as long as they wanted. When they were finished, they told the experimenter who then put the next object in the box. Overall, participants completed 27 trials (3 colors x 3 shapes x 3 views) in the first block.

In the second block all nine stimuli were placed in random order on the rotating disc and participants had to perform the single color adjustment task (motion match). Participants were instructed to watch at least one full rotation of the object (which took about 30 seconds).

### Analysis

Color settings were recorded in CIELab coordinates. For the analyses, we focus on the hue of the setting, i.e. the a* and b* coordinates, or the corresponding hue angle of the color settings. To determine whether the hue angle of the single color settings in the static condition varied as a function of paint, shape and view, we conducted a three-way ANOVA, and a corresponding two-way (shape x paint) ANOVA for the motion condition. To assess the hypothesized potential modulatory influence of object motion on single color appearance, we then performed a regression analysis, measuring the extent to which hue angles in the motion conditions could be predicted from hue angles in the static conditions, and whether hue angles tended to shift more towards values that were present near the contour regions of the object. Next, in order to identify what determines single color appearance specifically, we used regressions to predict the hue angle of single color settings in the static and motion conditions from the hue angle of the setting that had the maximum saturation, the most similar hue, and the highest coverage, as indicated by the observer.

## 3 Results & Discussion

### Influences of paint, shape and view on single color matches

Fig. 2 depicts participants’ color settings in CIE L*a*b* space for view 1, each type of paint (panels A-C), each shape (rows), and each task (column), as well as corresponding bar plots for single-multi- & motion conditions, where the perceived area of a given color on the object is indicated by the size of the bar segment.

**Figure 2.**
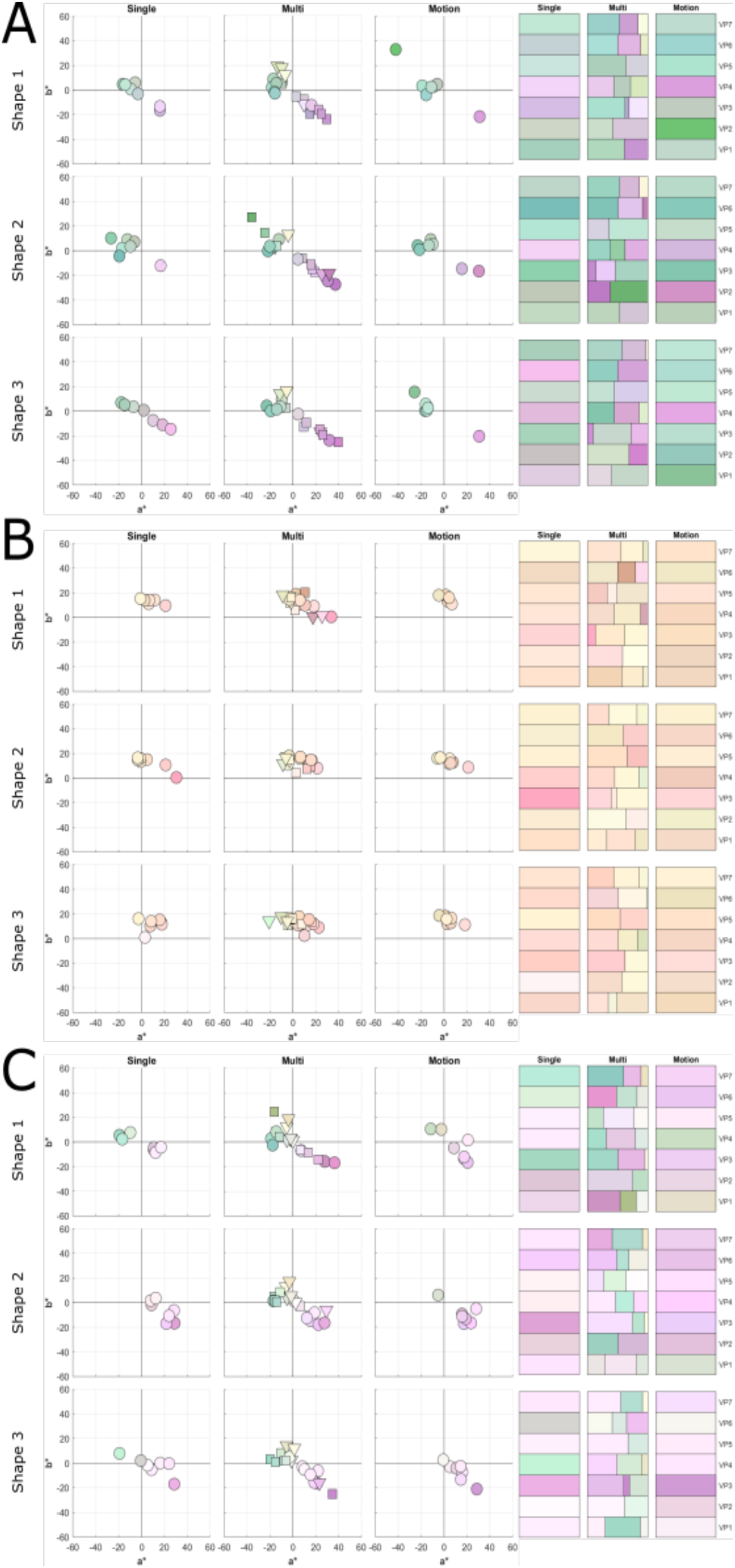
Color settings as a function of paint and shape complexity in the single and multi-match tasks. Shown are the results for static and motion conditions for view 1. On the left of each panel, color settings of observers in all tasks are plotted in CIELab. On the right, color settings are plotted as lists of horizontal bars for each observer, where the length of the bar segment for the multi setting task indicates the amount of coverage. For plotting, color values are converted from CIELab to RGB. Panels A-C show color settings for iridescent paints 1-3, respectively.

Overall, qualitatively, we can see that observers’ color settings varied not only with the type of iridescent paint (compare panels Fig2. A-C), but also across shapes (i.e. shape complexity), and views (also see Fig. 3). The three-way (paint x shape x view) ANOVA yielded a significant main effect of paint F(2, 162)=3.61 p<0.05, and a significant interaction of shape and view F(4, 162)=2.45 p<0.05. Following up the interaction with 3 one-way ANOVAs (one for each shape) we found, however, no significant effects of view, which makes this interaction challenging to interpret. A potential modulation of color setting by shape and view would be interesting. In Fig. 1 we see that the same kinds of colors exist on each shape and for each of the views, however, the amount and distribution of colored areas on the object’s surface vary. This change in color ‘layout’ or geometry could influence how we judge an object’s overall color. In future experiments we will investigate potential geometric effects in more detail.

**Figure 3.**
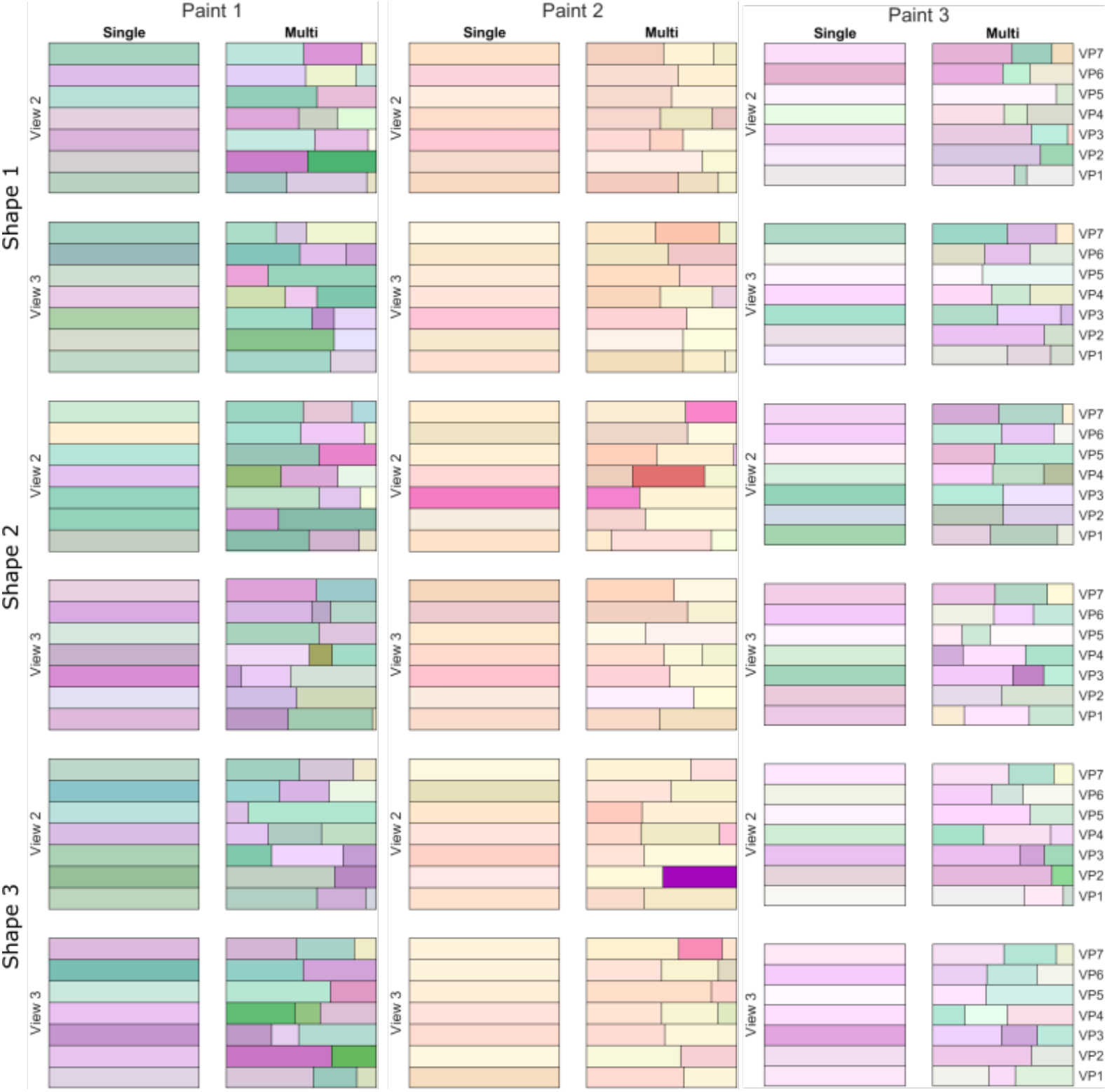
Color settings as a function of paint, shape and view. Shown are the color settings for single and multi-match tasks for views 2 and 3, for each of the three shapes and paints. Data are plotted as bars as in Fig.2 (right). Even though all three paints exhibit roughly a similar variety of hues, we find that overall, 3-color matches were more frequent for paints 1 (39 vs 24) and 3 (47 vs 16) but less frequent for paint 2 (29 vs. 34). This difference is likely due to the different degrees of hue similarity in each paint.

Interestingly, the significant interaction of view and shape to single color judgments disappears as the object rotates: in the motion condition, a two-way (paint x shape) ANOVA yields only a main effect of paint (F(2, 54)=12.87, p<0.05). This suggests that motion may have some ‘overriding’ effects over other factors such as shape and view. The rotating stimuli looked quite mesmerizing, with some of the colors swirling like large saturated highlights across the surface. Perhaps these effects captured the participan’t overall attention, or modulated their overall parsing of the stimuli, e.g. some kind of scision [19] separating the colors into base, i.e. belonging to the object (e.g. stationary, or slow moving parts) and highlight i.e, not belonging to the object (e.g. changing, or fast moving colors). We will further assess the changes in appearance in the motion condition below.

The variability in hue angle setting (of single match tasks) between observers was overall lowest for paint 2 (23.3) and about equal for paints 1 and 3 (53 & 50.72, respectively (Fig 4.). This is not surprising, since the hues present in paint 2 were located much closer in CIELab space than those for paints 1 and 3 (see also Fig. 1). A sign test comparing variability differences between static and motion conditions yielded a significant result z=2.5, p=.0123 (two tailed). Surprisingly, variability tended to decrease in the motion condition for the majority of paint, shape, averaged across views (8 out of 9). This reduction in variability would suggest that motion somehow yields visual information that can be utilized in a more similar fashion by observers. To understand what information these temporal changes contain is the goal of future work.

**Figure 4.**
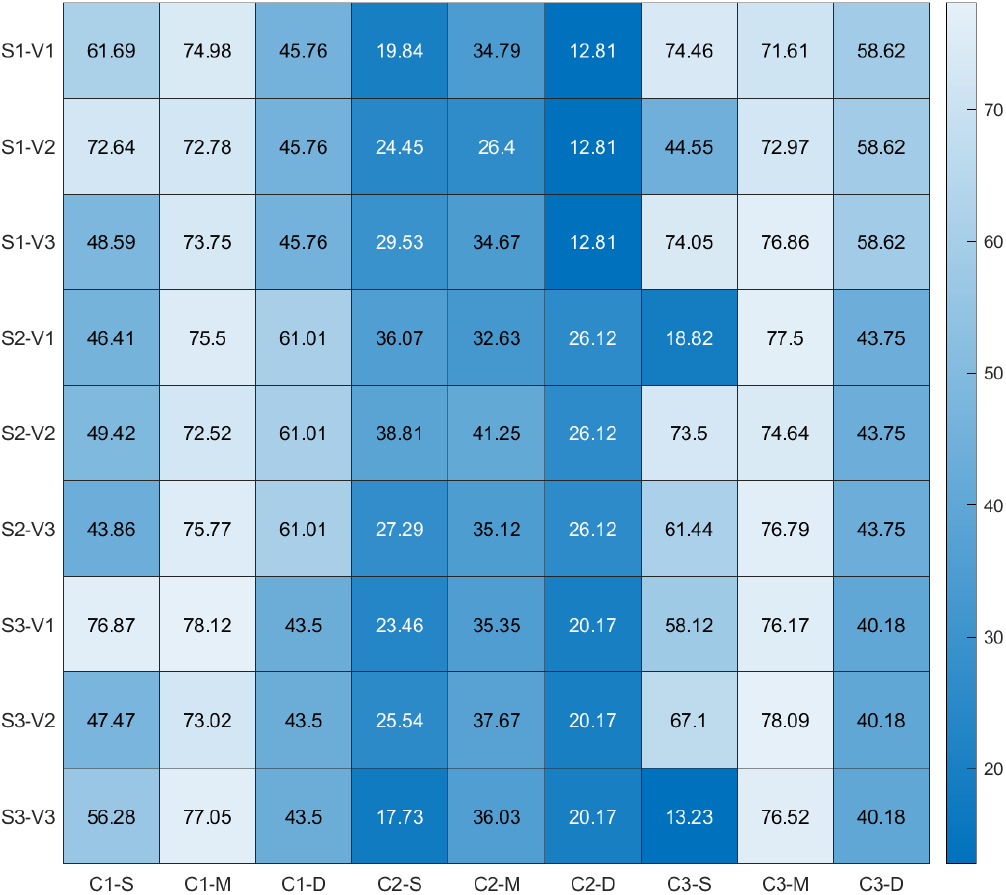
Mean observer variability in hue angle. Plotted is variability in hue angle as a function of shape (S1-3), paint (C1-3), view (V1-3) and task (S single, M multi, D motion). Darker colors imply less variability. Overall variability was lowest for paint 2.

### Modulatory effects of motion in single color matches

We see in Fig. 2, that single color settings for the same observer tended to change somewhat between static and motion conditions. Results of the regression analysis between hue angle settings of static and motion conditions are depicted in Fig. 5. Hue angle in the static condition was not a perfect predictor of hue angle in the motion condition (R^2^ =.17, R^2^ =.32, R^2^ =.03, for views 1, 2, and 3 respectively). Why might this be? For the simplest shape in Fig. 1, we see that central and peripheral regions on the object have distinct color appearances. However, this relation (center-edge color) gets more complicated as the shape gets more complex. When we rotated our iridescent objects, the central region tended to look more dynamic, almost like a colored highlight sliding across the object. If this was also what observers perceived, we expected their color matches to shift more towards the color that was present near the object boundary. However, we also expected this effect to vary with shape complexity and paint. To assess this, we performed k-means clustering on hue angles for single color settings, forcing the number of clusters to n=2, and counted the number of items in each cluster. We used the hue of the centroids obtained from settings of the simplest, sphere-like shape to label a cluster as ‘center’ or ‘edge’, e.g. for paint 1 purple is the center and green is the edge. And finally compare the counts in center and edge centroids across static and motion conditions. Fig. 6 shows that only for paint 1, the shift in hue angle occurred in the expected direction. For paint 2, the shift occurred in the opposite direction and for paint 3, motion did not modulate the number of settings corresponding to center and edge colors. A three-way ANOVA (paint x shape x rotating (yes, no)) supports these observations, yielding a significant main effect of paint F(2, 234)=12.56, p < 0.05 and a significant interaction of paint and rotating F(2, 234)=4.5, p < 0.05. Following up the interaction, we see that only for paint 1 and 2 hue angle settings tended to differ, which suggest that motion modulation depends on the similarity of hues present on an iridescent object (paints 1 a 2 exhibit colors that are more dissimilar in hue). Overall, while these effects of object motion on perceived color may be somewhat surprising, for other kinds of perceived surface material properties, like glossiness, its modulatory effects are well established [20].

**Figure 5.**
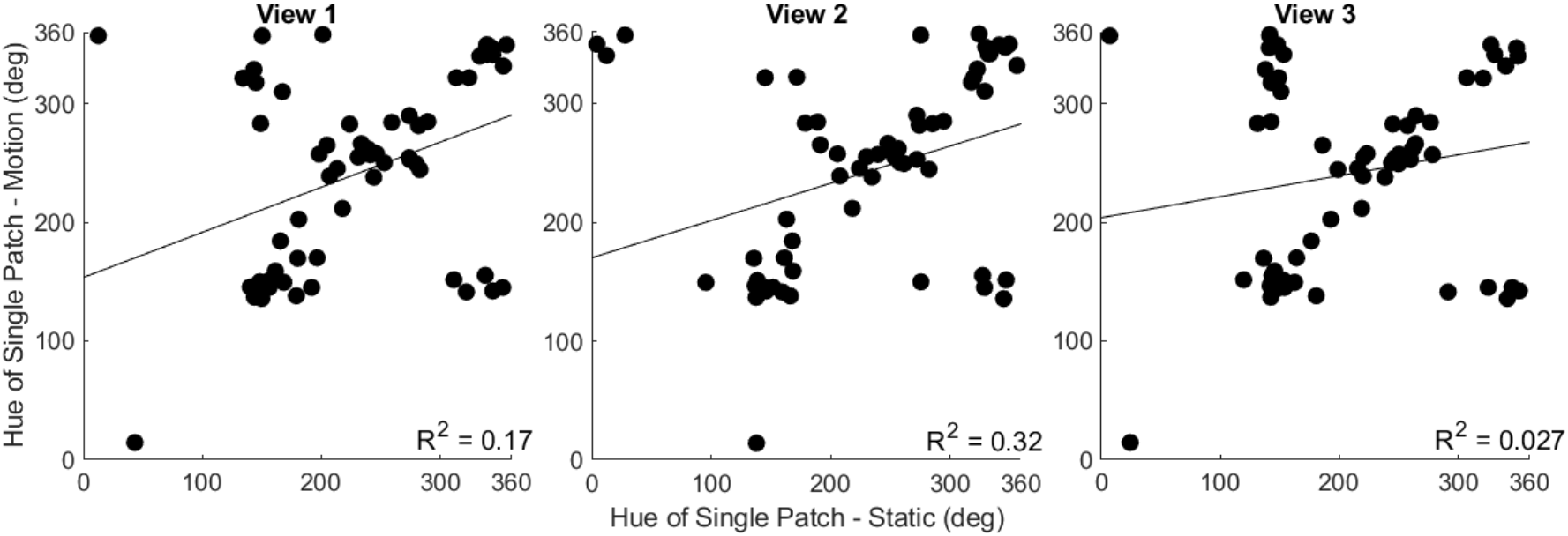
Predicting hue angle settings in the motion condition from hue angles in static condition. Shown are the scatter plots between the static single setting conditions (each view) and the motion condition, and corresponding regression lines. R^2^ denote the squared circular correlation coefficients.

**Figure 6.**
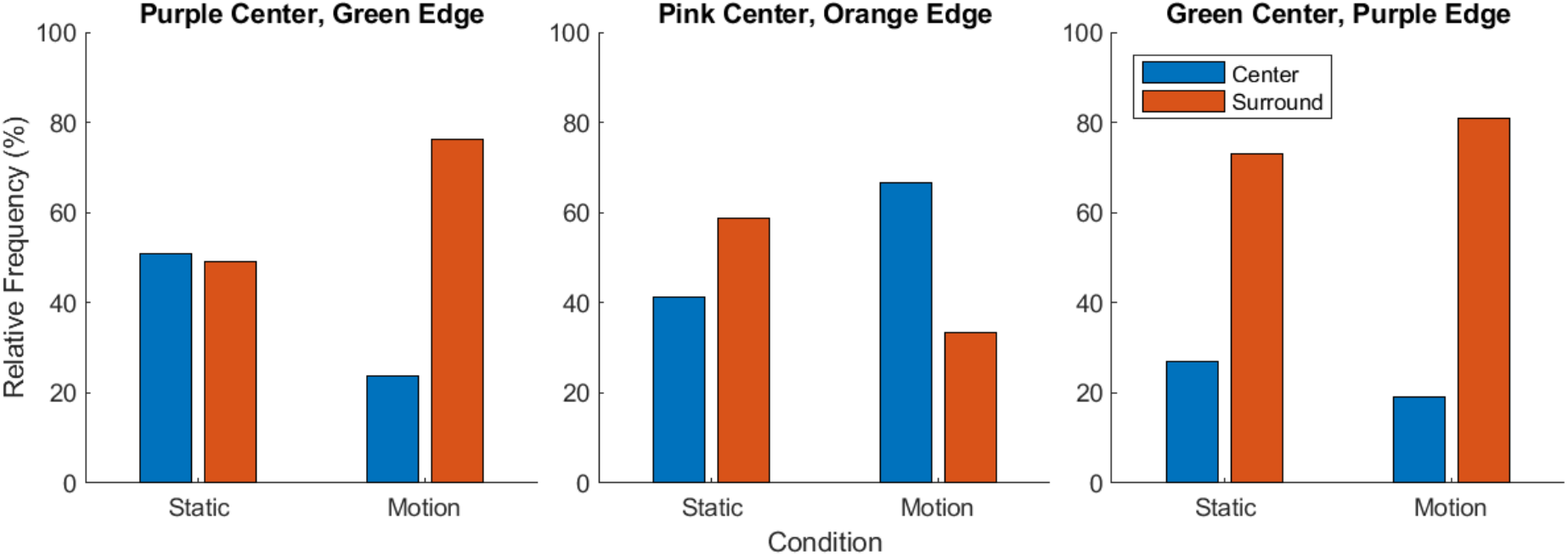
Center-to-edge shift of perceived hue between single static and motion condition. Y axis plots the relative frequency of settings being classified as center or edge color using k-means clustering. Panels 1-3 show results for corresponding paints. Results for the static condition combine data from all three views. See main text for details.

### Multicolor matches

The multi-color setting condition provided data for testing specific hypotheses of what perceived stimulus characteristics determine single color appearance. Observers were given the option of matching up to three prominent colors that they saw on the object, and to indicate how much the object was covered by the color they indicated. To identify what determines single color appearance, we predicted the hue angle of the single patch setting from the hue angle of the multi-color setting that had the maximum saturation, the most similar hue, and the highest coverage. Fig. 7 shows that overall, hue similarity was the best predictor. However, its strength varied across shape complexity and paints: on average it tended to be strongest for paint 1 (mean R^2^: .79) and weakest for paint 3 (mean R^2^: .2), strongest for shape 2 (mean R^2^: .67) and weakest for shape 1 (mean R^2^: .38). Also prediction strengths of single setting hue angle by max saturation and coverage varied across shape and paints, yet overall their prediction tended to be much weaker compared to hue similarity. In fact, for 6 out of the 9 shape x paint combinations, the hue similarity was the only significant predictor (i.e., 95% confidence interval of weight did not include 0). Only in two cases, maximum coverage and in just one case, highest saturation also contributed. Yet in all of these cases, the weight of hue similarity was always at least 3x as large, sometimes being 6x as large.

**Figure 7.**
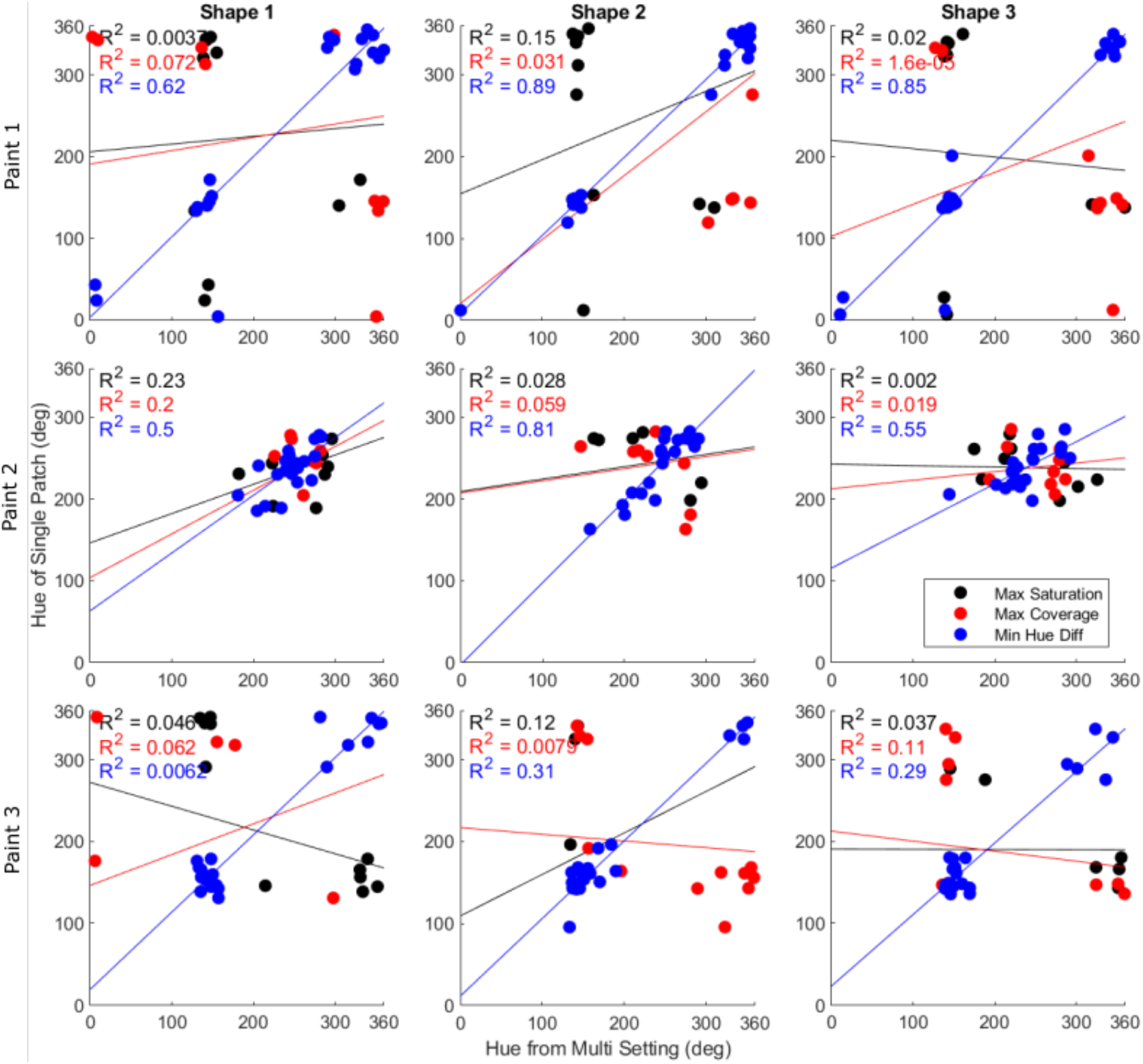
Predicting single setting hue angle from multi-color matches. For each paint and shape combination, we tested three potential strategies to determine the hue angle of the single setting color match: the max saturation in the multi-match condition (black regression lines and symbols), the most similar hue, i.e. minimum hue difference (blue symbols and regression lines), and the largest coverage (red symbols and regression lines). R^2^ is the circular correlation coefficient squared. All regressions with hue similarity as predictor were significant, with p< 0.0001.

To test whether the relative contributions of these three predictors change when observers judge a rotating object, we next repeated the regression analyses for settings obtained from the motion condition. Fig. 8 shows that overall, though with some exceptions, the strength of hue similarity as a predictor for hue angle decreased (R^2^ decreased on average by .13). The same pattern was true for the other two predictors. This may suggest that observers change their overall strategy for single color estimation when the object moves, perhaps they tend to mix colors or do some sort of temporal weighting of colors when the object is in motion [21][22], a possibility that we plan to test in future research.

**Figure 8.**
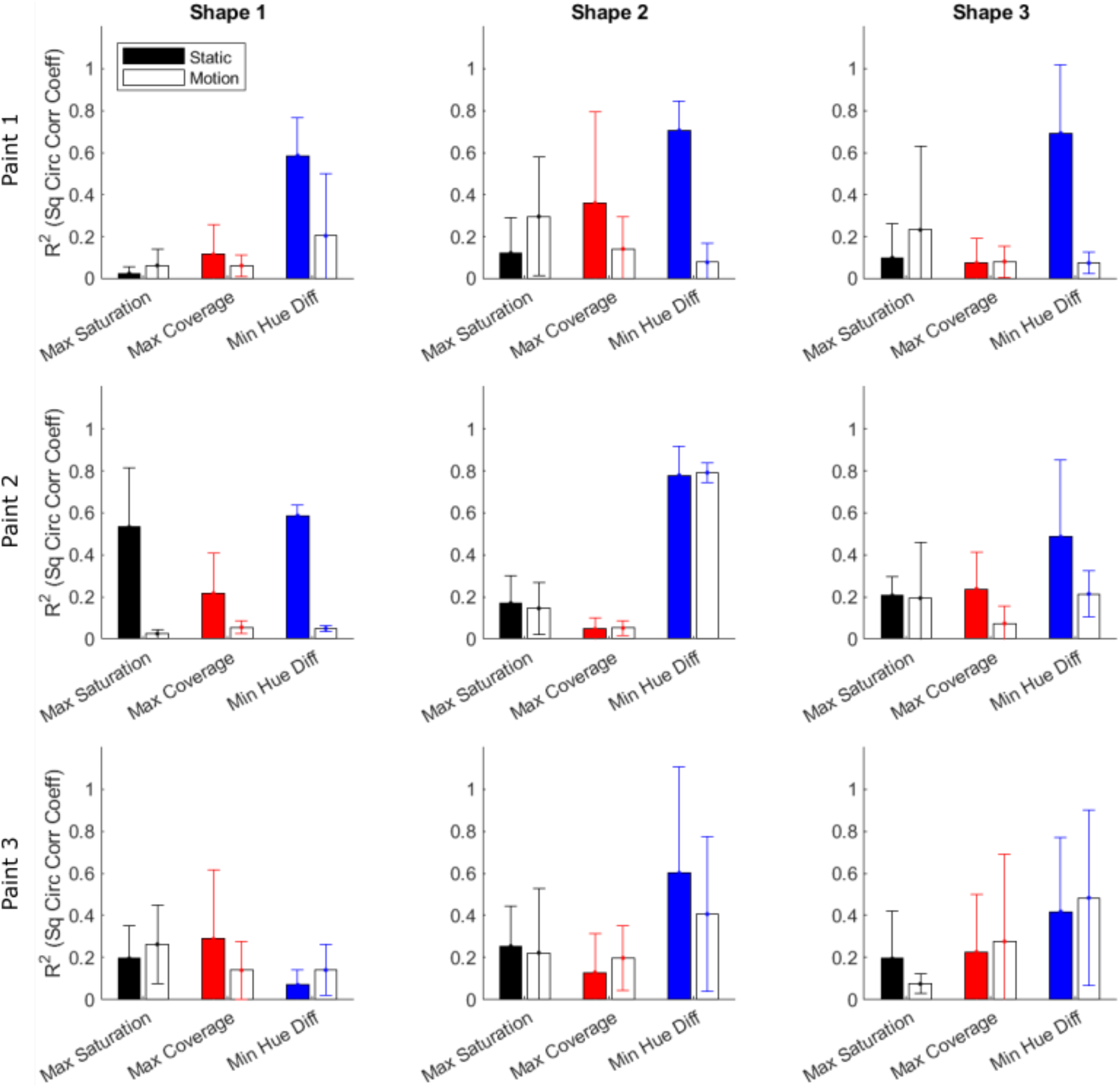
Changes in predictor strength between static and motion conditions. Plotted are the predictor strengths as bars for static (filled) and motion (empty) conditions as a function of paint (rows) and shape complexity (columns). Colors denote the three different predictors, as in Fig. 7.

## 4 Conclusion

We show that the single color appearance of iridescent real, three-dimensional objects is modulated by chromatic factors, spatial-relations and the characteristic dynamics of color changes that are typical for this type of material. In future studies, we will combine perceptual and physical measurements of the iridescent stimuli to fully understand how these factors interact.

